# A syndecan-based genetic approach to coat the surface of small extracellular vesicles with Nanobodies

**DOI:** 10.64898/2026.01.29.702589

**Authors:** Lukas Hyka, Sofie Meeussen, Rania Ghossoub, Michiel De Coster, Elke De Bruyne, Guido David, Patrick Chames, Pascale Zimmermann

**Affiliations:** Laboratory for Extracellular Vesicle Research, Department of Human Genetics, Katholieke Universiteit Leuven, Leuven, Belgium; Centre de Recherche en Cancérologie de Marseille (CRCM), Equipe Labellisée Ligue 2018, CNRS, Inserm, Institut Paoli Calmettes, Aix-Marseille Université, Marseille, France; Translational Oncology Research Center (TORC), Team Hematology and Immunology (HEIM), Vrije Universiteit Brussel, Brussels, Belgium; Centre de Recherche en Cancérologie de Marseille (CRCM) CNRS, Inserm, Institut Paoli Calmettes, Aix-Marseille Université, Marseille, France

**Author notes:** **Corresponding author:** Pascale Zimmermann.

**Keywords:** small extracellular vesicles, genetic engineering, Nanobody, Syndecan, EV-surface coating

## Abstract

Small extracellular vesicles (sEVs) are promising vehicles for targeted therapeutic delivery, but strategies for their surface functionalization remain limited. Here, we present a reliable and simple genetic approach that enables customized modification of sEV surfaces and supports enhanced sEV uptake by recipient cells. This strategy is based on the fusion of targeting moieties to the C-terminal fragment of syndecan-1 (SDC1-CTF), a peptide naturally enriched in sEVs. Combining various analytical approaches including single-vesicle analysis, we establish that this strategy enables decoration of up to 20% of secreted sEVs with nanobodies (Nbs). In quantitative bioluminescence assays, using concentrated conditioned media, we demonstrate that sEV-coating with anti-EGFR Nb supports enhanced sEV uptake by EGFR-expressing cells. This new strategy thus offers a robust and modular solution for endowing sEV surfaces with defined targeting properties to support further sEV-based therapeutic applications.

## Introduction

Extracellular vesicles (EVs) are membrane-limited organelles released by virtually any type of cell. EVs are heterogeneous in size but can be separated into small, typically 50-150 nm in diameter EVs (sEVs), and larger, typically 200-1000 nm in diameter EVs (lEVs), by different fractionation approaches. EVs are also heterogeneous in composition. They have been shown to carry multiple different proteins, RNAs and lipids and to be capable to deliver these molecules to cells, altering recipient cells (Valadi et al. 2007; Dixson et al. 2023; Doyle and Wang 2019). As EVs are considered non-immunogenic (Zhu et al. 2017), they are investigated for their use as delivery vehicles. In this context, the benefits of native EVs, as well as modified EVs are under intensive investigation. Native mesenchymal stromal cell-derived EVs have been shown to have anti-inflammatory and pro-angiogenic properties and are currently in clinical trials, for example in the treatment of chronic kidney diseases (Chen et al. 2020). Modified EVs include EVs presenting a surface molecule and/or containing bio-active or therapeutic cargo (Murphy et al. 2019). Advantageously, EVs can be modified to increase their circulation time (Du et al. 2021), orient their targeting (Jayasinghe et al. 2022; Alvarez-Erviti et al. 2011), increase their uptake (Noguchi et al. 2021) and/or deliver specific cargo (Dooley et al. 2021). Currently, modified EVs delivering their siRNA cargo targeting mutated K-Ras reached clinical trials for the treatment of pancreatic cancer (Mendt et al. 2018). Despite these advances the modified EV-based therapies are still limited by the low efficiency of cargo loading and by the large quantities of EVs necessary to observe the desired effects (Tang et al. 2019; Kennedy et al. 2021; Wiklander et al. 2015).

Here, we extend on prior work of our team indicating that syndecans (SDCs), interacting directly with syntenin, are implicated in sEV biogenesis (Baietti et al. 2012). One member of this family is SDC1, which is highly expressed in epithelial and plasma cells (Lambaerts et al. 2009). In particular, we showed that the proteolytically cleaved and membrane-associated C-terminal fragment (CTF) of SDC1 composes a major constituent of sEVs compared to its non-cleaved form (Baietti et al. 2012). Later we observed that in epithelial cells, like MCF-7 cells, SDC1-CTF is better than tetraspanins at predicting sEV numbers (Castro-Cruz et al. 2023). SDC1-CTF has been shown to serve as a sorting mediator to coat EVs with decoy moieties (Gupta et al. 2021). Here, we (i) evaluated whether fusing SDC1-CTF to targeting moieties, such as Nanobodies (Nbs), via a protease-resistant linker, would result in customized surface coating of sEVs and (ii) tested whether such coating would help sEV-uptake (binding and/or internalization) by target cells. Nbs are single-domain antibodies of small size (15 kDa), corresponding to the variable domain of heavy chain-only antibodies that are circulating in Camelidae. They are characterized by high stability, specificity and low immunogenicity (Even-Desrumeaux et al. 2012).

## Materials and methods

### Reagents & antibodies

Fetal bovine serum (FBS, Sigma, 1681067), DMEM (Gibco, 41965039), XtremeGene 9 (Roche, 6365809001), Geneticin (Gibco, 10131027), EV-depleted FBS (FBS depleted of intrinsic EVs by overnight (18 h) centrifugation at 100,000 x g, followed by filtration through sterile 0.45 µm filters (Merck), Amicon 10 kDa filters (Merck, UFC901024), and Furimazine (DC chemicals, DC70371) reagents were used. The anti-SDC1-CTF and syntenin-1 antibodies are homemade and were described before (Lories et al. 1992). The antibodies against CD9, CD63 were a kind gift from Prof. Eric Rubinstein (Charrin et al. 2001) (Université Paris-Sud, Inserm UMRS 935, Villejuif, France). Calnexin (Abcam, Ab10286), myc (CellSignaling, #2276), EGFR extracellular domain (Santa Cruz, sc-120) and Nanoluciferase (Promega, N7000) antibodies and directly conjugated (Alexa 647 and Alexa 488) anti-Variable Heavy domain of Heavy chain antibody (VHH) antibodies (Jackson ImmunoResearch, 128-605-230 and 128-545-230) were used per manufacturer recommendations.

### Cell culture and transfections

HEK293, and Panc-1 cells (ATCC) were cultured in DMEM supplemented with 10% FBS. For transfections, HEK293 cells were plated the day before so to achieve ∼40% confluency at the time of transfection. Cells were transfected with expression vectors using XtremeGene 9 according to the manufacturer’s protocol. Following transfection, cells were cultured for 48 hours before the start of the conditioning of the medium for EV enrichment either in DMEM supplemented with 10% EV-depleted FBS for the isolation by differential ultracentrifugation or size-exclusion chromatography or in serum-free DMEM for the isolation by concentration (also see below).

### Plasmid generation

Myc-tagged SDC1-CTF, the platelet-derived growth factor receptor transmembrane domain (PDGFR-TM) and Prostaglandin F2 receptor negative regulator (PTGFRN) myc-tagged anti-Nef chimeras and anti-Nef, anti-EGFR, non myc-tagged SDC1-CTF chimeras were ordered as gBlocks (IDT DNA) and inserted into pcDNA3 plasmid by Gateway cloning. Plasmid encoding Nanoluciferase was kindly provided by prof. Clotilde Thery (Institut Curie, Paris). The Nanoluciferase fragment was fused to syntenin by overlapping PCR and Gateway recombination cloning. Amino acid sequences of prepared constructs are listed in Table S1.

### Preparation of stable clones

Cells were transfected as described above with linearized expression vectors. 48 hours after transfection, cells were put under Geneticin selection at 300 µg/ml or 800 µg/ml for HEK293 and Panc-1 respectively. These concentrations were determined as the minimal doses inducing full cell death after 14 days. Resistant colonies were picked after 21 days and expanded separately. Proper transgene expression was determined by Western blot with an anti-SDC1 antibody directed against the intracellular domain. Transgene expression in total clonal population (homogeneity) was confirmed by confocal microscopy after immunostaining with the same antibody.

### Immunofluorescence analyses

The cells were cultured on eight-well chamber slides (Nalgene Nunc, 177402) overnight, fixed with 4% PFA for 15 min, washed with PBS, and permeabilized with 0.05% saponin in PBS for 20 min, in the presence of 0.3% BSA for blocking non-specific reactions. After incubation with primary antibodies, cells were incubated with Alexa-conjugated secondary antibodies (Invitrogen). Nuclei were stained using DAPI and the samples were mounted with ProLong Glass Antifade Mountant (ThermoFisher, P36982). Samples were observed with an Olympus Fluoview 1000 confocal microscope. Images were analyzed using ImageJ/Fiji (National Institute of Health, Bethesda, MD) and Photoshop (Adobe, San Jose, CA) software.

### Preparation of conditioned media and EV enrichment

Transfected cells were washed once with fresh culture medium (DMEM) before media were conditioned for 16 hours (5 ml medium per 10 cm plate of 1.2 million cells (day 1)). DMEM supplemented with 10% EV-depleted FBS was used when EVs were enriched by differential ultracentrifugation (dUC) or size exclusion chromatography (SEC). DMEM without FBS was used when EVs were enriched by concentration.

#### Differential ultracentrifugation (dUC)

Conditioned media were subjected to three sequential centrifugation steps at 4°C: 10 min at 1,500 x g (50 ml Falcon tube, rotor Sigma 12310), to remove cells and cell debris; 30 min at 10,000 x g (10K; 50 ml Falcon tube, rotor Sigma 12310) to pellet large particles-EVs (lEVs); and 1 h at 100,000 x g (100K; Beckman Open-Top Thinwall Ultra-clear Tube rotor SW32Ti) to pellet small particles-EVs (sEVs). 100K pellets were washed in PBS, centrifugated for 1 h at 100,000 x g (Beckman Microfuge Tube Polyallomer, rotor TLA-55), and re-suspended in 40 µl of PBS, as were 10K pellets for further analysis by Western blot. The corresponding cell layers were kept on ice, scraped in lysis buffer (Tris-HCl 25 mM, pH 7.4, NaCl 150 mM, EDTA 1 mM, NP40 1% and protease inhibitors), and incubated for 45 min at 4°C under gentle agitation. Cell lysates were cleared by centrifugation at 10, 000 x g at 4°C for 10 min before analysis by Western blot.

#### Size exclusion chromatography (SEC)

Conditioned media were processed as for dUC for cell debris and lEVs removal. The 10K pellet supernatant was loaded on Amicon 10 kDa filters and concentrated by 2, 000 x g centrifugation until the volume dropped from ∼10 ml to ∼500 µl (20 x concentration). The concentrated medium was loaded on qEV original 35 nm column (Izon), and 12 fractions of 0.5 ml were collected. The fractions were centrifuged at 10, 000 x g, 4°C for 40 min on Amicon 3 kDa filters to concentrate the volume to ∼80 ul. Aliquots (20 ul) of these samples were characterized by Western blot.

#### Preparation of concentrated conditioned medium (CCM) sEVs

Conditioned media were processed as for dUC for cell debris and lEVs removal. The 10K pellet supernatant was loaded on Amicon 10 kDa filters and concentrated by centrifugation at 2, 000 x g until the volume dropped from ∼10 ml to ∼300 µl (33 x concentration). This concentrated supernatant was referred to as the CCM sEVs fraction and was characterized by Western blot or used in uptake experiments.

### Western Blotting

Cleared cell lysates and EV-enriched fractions were resuspended in PBS, boiled for 5 min in Laemmli buffer (8.29 % glycerol, 1.90% SDS, 9.52 mM Tris, 0.95 mM EDTA, bromophenol blue) and fractionated by SDS-PAGE (NuPAGE 4-12% Bis-Tris gels, Invitrogen) before transfer to nitrocellulose membranes (Hybond-C extra, 0.45 µm, GE Healthcare). Cell lysates loaded always corresponded to approximately 20 000 plated cells (day 1) – equivalent to 20 µg protein according to bicinchoninic acid assay. EV-enriched fractions were most often collected from 3.6 million plated cells (day 1), unless specified differently. Membranes were probed with primary and HRP-conjugated secondary antibodies, followed by chemiluminescence detection (Perkin Elmer). Band intensities were quantified using ImageJ/Fiji.

### EV uptake experiments

CCM sEVs (for preparation see above) of HEK293 clones stably expressing SDC1-CTF chimera and transiently expressing Nanoluciferase-syntenin (NL-synt) were incubated with Panc-1 recipient cells (plated for 16 h and washed before the start of the experiment with DMEM without FBS) at different concentrations, for different periods of time, as indicated. CCM sEV doses administered to recipient cells were determined by the number of plated producing HEK293 cells to the number of plated recipient Panc-1 cells, expressed as ratio of producing:recipient (p/r). Cells were washed twice with versene (Gibco) and incubated with Furimazine at 8.3 µM for 5 min before the luminescence measurement on a Victor Nivo plate reader (PerkinElmer).

### Measurement of particle concentration and size

EV-enriched dUC fractions obtained from equal cell amounts after identical culture durations were analyzed by nanoparticle tracking analysis using the Twin ZetaView^®^ instrument (Particle Matrix) or Microfluidic Resistive Pulse Sensing (MRPS) using the nCS2 instrument (Spectradyne^®^). For ZetaView, samples were diluted in PBS and for each sample, particle size and concentration were measured at least four times at 22°C (laser 488 nm, sensitivity 68, shutter 100, frame rate 30). For nCS2, samples were diluted in 1% Tween-20 PBS (filtered with a 0.02 µm syringe), and 5 µL aliquots were loaded into C-400 microfluidic cartridges. Data acquisition and analysis were performed using the nCS2 Viewer software. Measurements with both instruments were corrected for dilution and background subtracted.

### Measurement of Nanobody positive EVs

The presence of Nanobodies on CCM sEVs was evaluated using three independent methods: direct stochastic reconstruction microscopy (dSTORM) using the Oxford NanoImaging microscope (ONI^®^), nano flow analysis (CytoFLEX Nano^®^) and MRPS-coupled fluorescence (ARC, Spectradyne^®^).

#### dSTORM-microscopy

CCM sEVs were stained overnight with Alexa 561-conjugated α-pan-tetraspanin antibodies (targeting CD63, CD81 and CD9) (ONI^®^) and Alexa 647-conjugated α-VHH antibodies. Stained CCM EVs were immobilized on biotinylated coverslips prepared as previously described (Margeat et al. 2006) and coated with phosphatidylserine capture reagent (ONI PS capture EV Profiler 2 kit) before imaging. Laser powers were set to 12% (640 nm) and 80% (560 nm). Images were analyzed using CODI software (ONI^®^). A particle with at least 4 dots (‘localizations’) in the 561 nm channel was classified as a single CCM sEV. Particles additionally containing at least 2 dots (‘localizations’) in the 647 nm channel were classified as Nb-positive sEVs.

#### CytoFLEX Nano^®^ flow analysis

CCM sEVs were incubated overnight with Alexa 488-conjugated α-VHH antibody, washed using Exo-spin™ (Cell Guidance Systems) to remove the antibody excess, and measured on a CytoFLEX Nano^®^ instrument (Beckman Coulter). Instrument calibration was performed using Megamix-Plus FSC and SSC beads (Biocytex, Marseille, France). Samples were run at 30 μL/min. The gating was set up on Mock CCM sEVs to remove the bulk of the dense signal (in blue) and was transposed to all the other samples. To compare with other single-EV approaches used in this study, gating was performed to the same antibody concentration (1:20). Data were analyzed with CytExpert Nano software (Beckman Coulter).

#### MRPS coupled fluorescence

CCM sEVs were stained as described for CytoFLEX Nano^®^ flow analysis except that removal of antibody excess could be omitted. Laser powers were set to 10mW for both lasers. The APD detector gains were set to 10 for all detectors.

### Statistical analysis

Statistical comparisons were performed using Student’s t-test or ANOVA followed by Tukey’s multiple comparison test. Graphs depict means ±SEM from at least three independent experiments, unless otherwise stated. Statistical analyses were conducted using GraphPad Prism v9.0.

## Results

### A SDC1-CTF-based approach for coating sEV surfaces

To coat the surface of sEVs, we first prepared a DNA construct comprising a synthetic open reading frame encoding (i) the signal peptide of SDC1 to ensure efficient membrane translocation and the correct topology of the chimera, (ii) a myc-tagged version of a Nb directed against the viral protein Nef (Bouchet et al. 2011), (iii) the juxta-membrane domain of CD4 (Fitzgerald et al. 2000) to promote the resistance of the chimera to proteolytic cleavage, and (iv) the membrane-spanning and cytosolic domain parts of SDC1-CTF expected to allow efficient sEV sorting (Fig. 1A). When examining cell lysates of HEK293 cells overexpressing these chimeric proteins by Western blot with antibodies directed against the SDC1 cytoplasmic domain, we observed a single band corresponding to the full-length chimera (Fig. S1). When examining secretomes fractionated by dUC, we detected two signals in the sEV-enriched fraction (100K pellet). One at the expected size for the full-length chimera and one corresponding to a cleavage product around the size of SDC1-CTF. The respective abundance of full-length and cleavage product was depending on the precise structure of the CD4 juxta-membrane peptide. Using a CD4 juxta-membrane peptide containing a proline (P) (Fig. 1A, red) instead of an arginine (R) used by others (Fitzgerald et al. 2000), supported the secretion of sEV strictly decorated with full-length chimera (Fig. S1). We therefore proceeded with constructs containing a proline.

**Figure 1.**
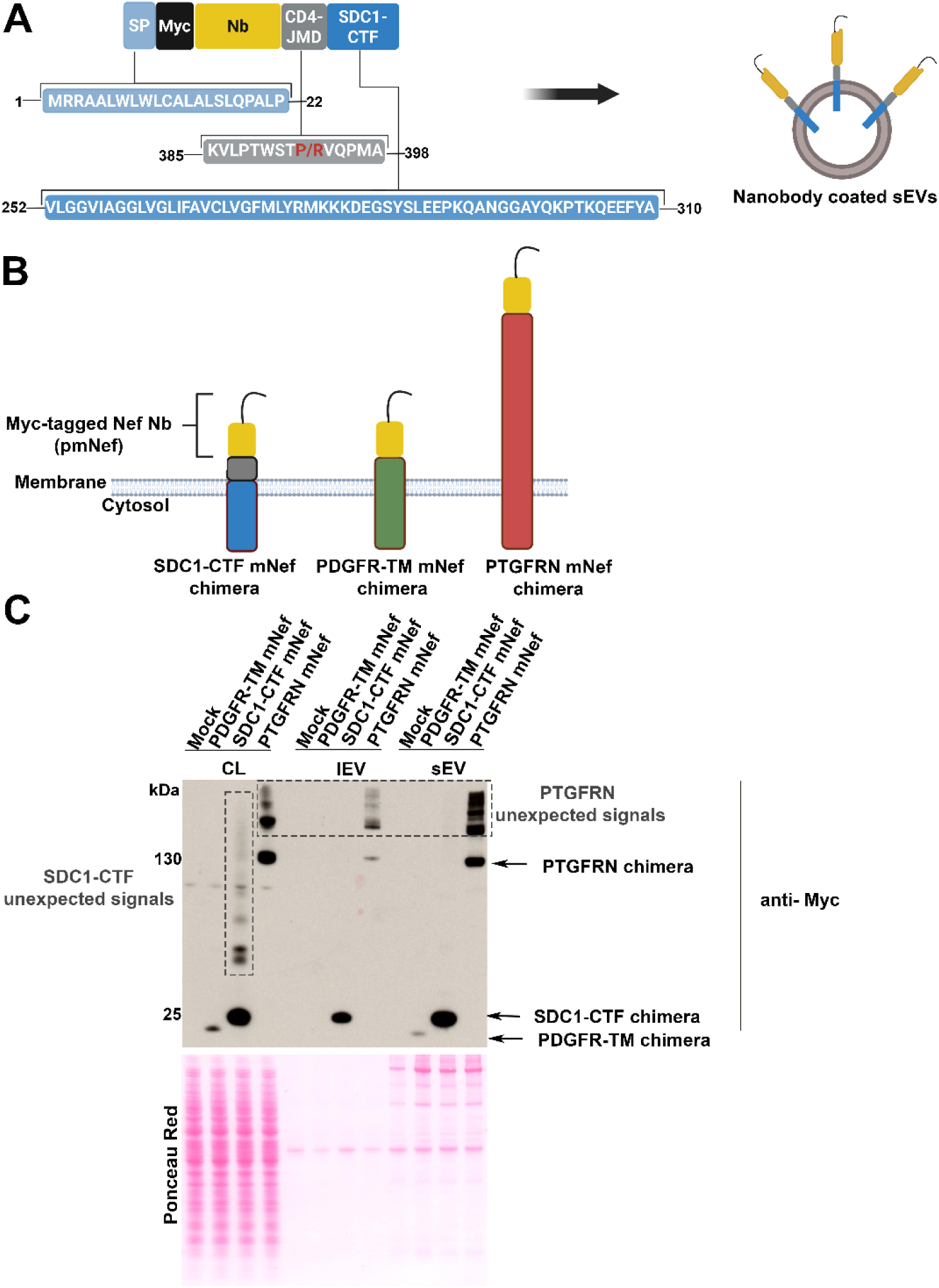
Comparison of various strategies for the coating of EVs with a Nanobody. **(A)** Schematic representation of the open reading frame (ORF) included in the pcDNA3.0 eukaryotic expression vector for the production of SDC1-CTF Nef chimera aimed at coating sEV surfaces. The ORF consists of five parts encoding: the SDC-1 signal peptide (light blue), the myc-epitope (black), the anti-Nef Nb (yellow), the CD4 juxtamembrane domain (CD4-JMD, grey), and the C-terminal fragment of syndecan-1 (SDC1-CTF, blue). Amino acid positions in full proteins are indicated. **(B)** Schematic representation of PDGFR-TM, SDC1-CTF and PTGFRN myc-tagged Nef Nb chimeras used in **C**. **(C)** Illustrative Western blot of cell lysates (CL) and extracellular vesicles (EVs) secreted by HEK293 cells transiently transfected with expression vectors encoding the different chimeras described under **B**. Mock refers to cells transfected with an empty vector, used as negative controls. EVs were enriched by differential ultracentrifugation (dUC). Large EV (lEV) and small EV (sEV) fractions correspond to the 10K and 100K pellets respectively. Blots were probed with anti-myc antibodies allowing comparison of the different chimeras. Ponceau red was used as loading and transfer control. CL correspond to 80.000 cells. Secretomes were collected from the conditioned media of 3.6 x 10^6^ cells. Note the presence of unexpected high-molecular-weight signals. Related to **Fig. S1** and **S2**.

We then compared our sEV-coating strategy with two previously described strategies. First, we used the transmembrane domain of the platelet-derived growth factor receptor commonly used for membrane anchoring (Zhang et al. 2014; 2025), and second the Prostaglandin F2 receptor negative regulator full-length receptor, a sorting moiety previously used for EV surface modification in the industry (Dooley et al. 2021). To enable comparison, we prepared myc-tagged Nef Nb-fusions, further referred to as SDC1-CTF mNef, PDGFR-TM mNef and PTGFRN mNef (Fig. 1B). We analyzed the lysates and secretomes of cells transiently transfected with these constructs. The ‘PDGFR-TM strategy’ turned out to be poorly effective in terms of obtaining Nb in EV fractions (Fig. 1C, Fig. S2). In contrast, the PTGFRN and SDC1-CTF strategies appeared roughly similar in supporting high Nb sEV sorting (Fig. 1C, Fig. S2). Nevertheless, we detected unexpected high molecular weight signals in lysates for SDC1-CTF transfectants (Fig. 1C, vertical dashed rectangle), and in lysates and EV fractions for PTGFRN transfectants (Fig. 1C, horizontal dashed rectangle).

To clarify whether the unexpected high-molecular weight signals may result from cellular stress associated with transient transfections, we prepared polyclonal cell populations stably overexpressing the SDC1-CTF and the PTGFRN, further referred to as SDC1-CTF pmNef and PTGFRN pmNef. When performing Western blot analysis of lysates and secretomes, we no longer observed unexpected signals for the SDC1-CTF pmNef, while the PTGFRN pmNef remained ‘problematic’ (horizontal dashed rectangle) – generating high molecular weight unexpected signals (Fig. 2A, Fig. S3). Western blots with EV markers like syntenin, CD9 and CD63 and the endoplasmic reticulum marker calnexin (supposedly absent from EVs) confirmed the quality of our EV-fractions. Of note, these blots also revealed that the SDC1-CTF pmNef does not promote lEV sorting, unlike the PTGFRN pmNef. When quantifying the respective sEV sorting efficiency (only taking into account signals corresponding to the expected sizes of the chimeras), we observed no significant difference between the SDC1-CTF and PTGFRN strategies (Fig. 2A right). Of note, Nano Particle Tracking analysis (NTA – ZetaView^®^) revealed no significant difference in particle concentration or median size between control and chimera secretomes (Fig. 2B).

**Figure 2.**
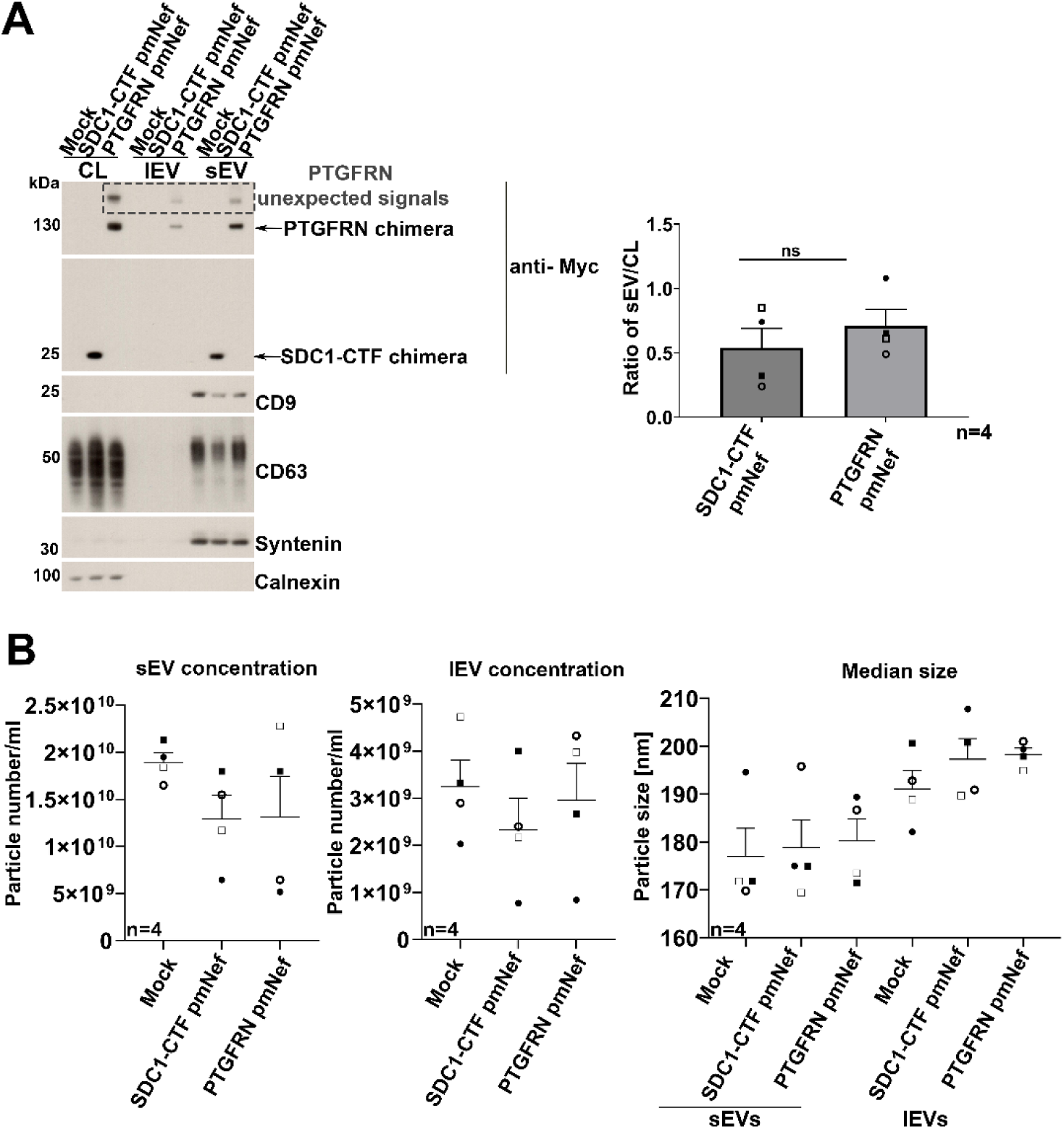
The SDC1-CTF chimera has a clean sEVs profile (compared to the PTGFRN chimera) and does not significantly influence EV concentration and size. **(A)** Left, illustrative Western blot of cell lysates (CL), large EVs (lEV) and small EVs (sEV) secreted by HEK293 cells stably expressing SDC1-CTF and PTGFRN Nef Nb chimeras, as indicated on top. Mock refers to cells transfected with an empty vector, used as negative controls. Blots were probed with anti-myc antibodies to allow comparison between the chimeras and other sEV markers (syntenin, CD63, CD9) and a negative control (Calnexin). CL correspond to 20.000 cells. EVs were collected from the conditioned media of 3.6 x 10^6^ cells. Note the presence of unexpected high-molecular-weight signals for the PTGFRN pmNef EVs and CL. Right, histogram illustrating that SDC1-CTF and PTGFRN chimeras are sorted to sEVs with similar efficiencies. Anti-myc signals obtained in Western blot for sEV fractions and CL were expressed as a mean of the sEV/CL ratio from 4 independent experiments, as indicated. Bars represent mean values + SEM. Student’s t-test was applied to assess statistical significance. **(B)** Graphs illustrating particle concentrations and their median size as detected in NTA measurements (ZetaView^®^) for sEV and lEV (obtained by dUC from equal cells amounts after identical culture durations), as indicated. Related to **Fig. S3.**

Taken together, these results indicate that SDC1-CTF and PTGFRN pmNef are equally efficient in sorting Nbs to sEVs. Yet, the SDC1-CTF pmNef offers more strict specificity for sEVs and poses no quality control concerns.

### Validation and quantification of Nanobody coating by single EV analytical approaches

Satisfied by these first datasets, we proceeded with the preparation of HEK293 clones homogeneously expressing the SDC1-CTF Nef chimera (Fig. 3A), this time omitting the N-terminal myc-epitope to avoid compromising Nb-activity. Using the highest expressing clone, further referred to as cNef, we validated the correct topology of the chimera. We therefore compared signals obtained with antibodies directed against SDC1 cytosolic domain (C-terminal) and directed against the Nef Nb (N-terminal) on permeabilized and non-permeabilized cells (Fig. 3B). The presence of the chimera in cNef sEVs was confirmed by dUC (Fig. S4A). No significant difference in particle concentration or median size between control and chimera sEVs was measured by Microfluidic Resistive Pulse Sensing (MRPS) using the nCS2 instrument (nCS2, Spectradyne^®^) (Fig. 3C). Conditioned media were also analyzed by size exclusion chromatography (SEC). We documented that the chimera strictly co-elutes with the sEV markers syntenin, CD63 and CD9 - before the protein rich-fractions highlighted by Ponceau Red (Fig. 3D, Fig. S4B). To determine the fraction of sEVs that are coated with Nbs, we cultured the HEK293 cells stably expressing the SDC1-CTF cNef chimera and Mock-transfected controls in the absence of serum, cleared the conditioned media of cell debris and 10K particles, and concentrated it by low-speed centrifugation. This concentrated conditioned medium sEVs fraction is referred to as CCM sEVs (Fig. 4A). We proceeded with the analysis of CCM sEVs using directly labelled anti-Nb antibodies by various single EV analytical approaches. First, we used nano flow analysis. After determining an optimal antibody concentration (1:20), we could estimate, by subtracting the signal obtained with Mock CCM sEVs that ∼ 6% of the anti-Nef CCM sEVs particles are Nb-coated (Fig. S5). Second, we used a super-resolution microscopy approach (ONI^®^). In this approach the anti-Nef CCM sEVs were labelled with a pan-tetraspanin antibody (CD9, CD63, CD81) and anti-Nb antibodies coupled to another dye (Fig. 4B). From 3 independent biological repeats (Table S2), we observed a mean Nb positivity of 30% (+/- 2% SEM) on the anti-Nef CCM sEVs. Yet, CCM sEVs of Mock-transfected controls gave a mean Nb positivity of 10% (+/- 2% SEM). We thus concluded that approximately 20% of the tetraspanin-positive sEVs contain anti-Nef Nb. Third, we used an MRPS fluorescence-coupled instrument (ARC, Spectradyne^®^) and measured from 3 independent experiments a mean of ∼10% (+/- 1% SEM) Nb-coated particles (Fig. 4C).

**Figure 3.**
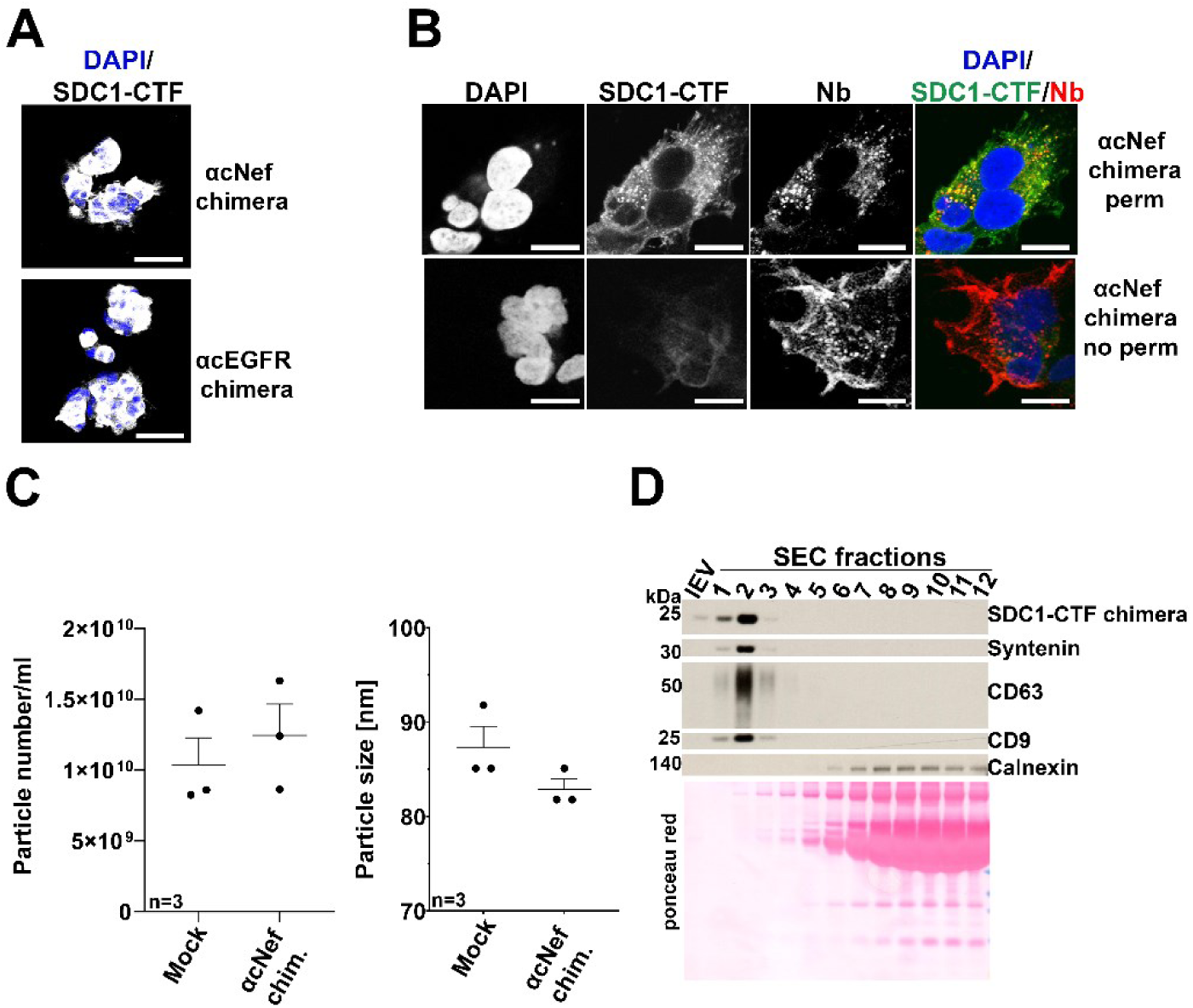
Characterization of the small EVs secreted by a HEK293 clone stably and homogeneously expressing SDC1-CTF chimeras. **(A)** Confocal micrographs of HEK293 clones stably expressing SDC1-CTF chimeras with anti-Nef (cNef) or anti-EGFR (cEGFR) Nbs, as indicated. Cells were stained with an antibody directed against the intracellular domain of SDC1 (grey). Nuclei were stained with DAPI (blue). Scale bar is 25 µm. Note the homogenous expressions of the chimeras. **(B)** Confocal micrographs of cNef HEK293 cells permeabilized (upper part) or not (lower part) before staining with an antibody directed against the intracellular domain of SDC1 (green in merge) and anti-Nb antibodies (red in merge) directed against the extracellular part of the chimera. Nuclei are stained with DAPI (blue in merge). Scale bar is 25 µm. Note the absence of SDC1 staining in non-permeabilized cells, indicating that the topology (outside-out, inside-in) of the construct is correct. The decrease of nanobody signal after permeabilization likely reflects partial removal of proteins during the IF procedure. **(C)** MRPS measurements illustrating particle concentration (left) and size (middle). sEVs were obtained by dUC. Mock refers to EVs from empty vector transfected cells. **(D)** Illustrative Western blot analysis of lEVs (first lane) and the conditioned medium fractionated by size-exclusion chromatography (SEC). Signals for the chimera and various EV marker proteins are indicated on the right. Mock refers to cells stably transfected with an empty vector. Ponceau red was used as loading and transfer control. Note that the chimera is solely present in the SEC-derived fractions positive for other sEV markers. Related to **Fig. S4**.

**Figure 4.**
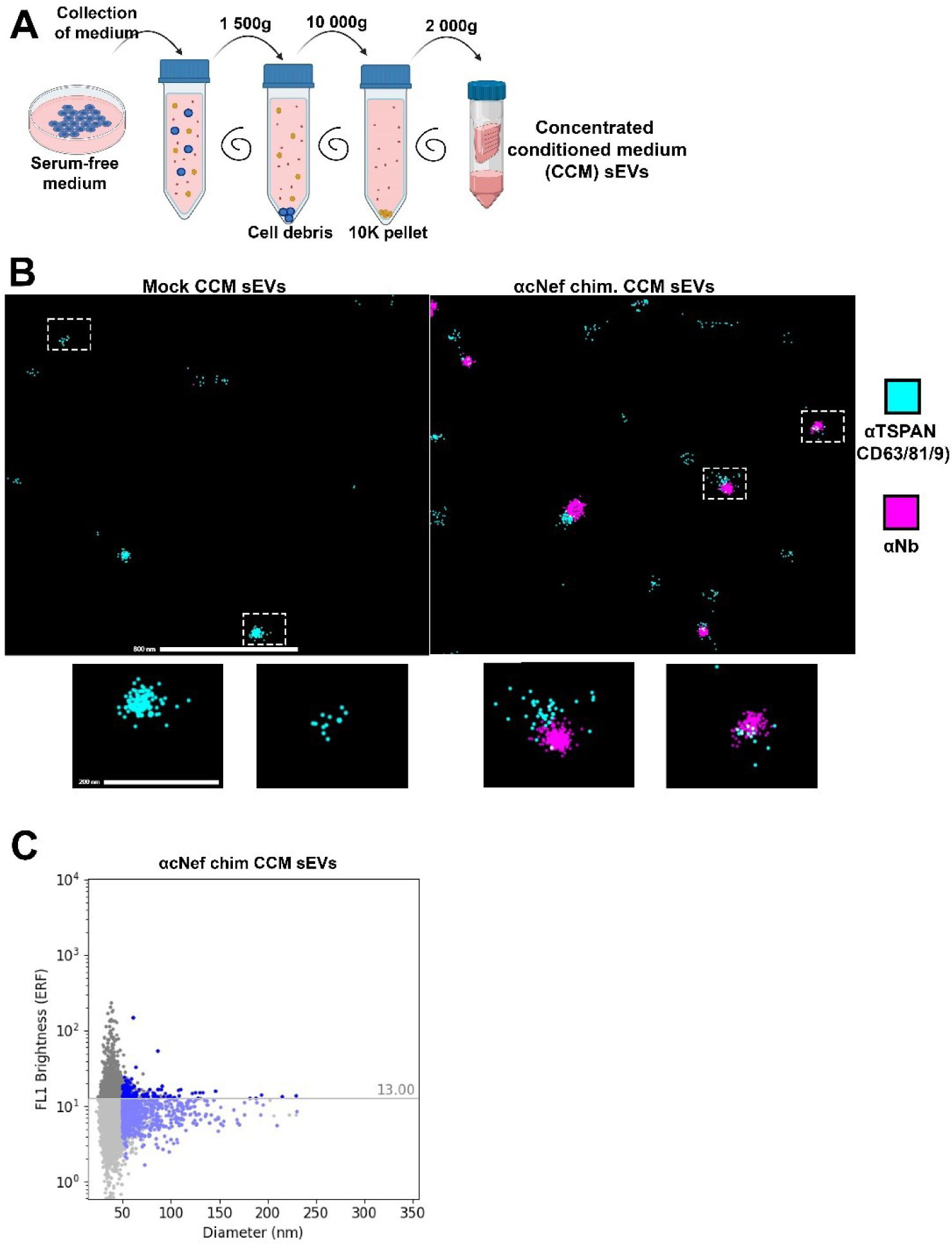
Single EV characterization of the small EVs in concentrated conditioned media (CCM sEVs) secreted by cNef HEK293. **(A)** Schematic representation of the procedure used to concentrate serum-free conditioned media to obtain concentrated conditioned medium (CCM) sEVs. **(B)** Representative micrographs obtained for Mock (left) or cNef (right) CCM sEVs by ONI^®^ microscopy. Anti-tetraspanin (CD9, CD63, CD81) signals are in cyan while anti-Nb signals are in magenta. Scale bars correspond to 800 nm for field view and 200 nm for inserts. **(C)** Illustrative representation of MRPS measurements with cNef CCM sEVs. Y-axis represents brightness intensity. X-axis represents particle diameter. Related to **Fig.S5**.

Taken together, the data indicate SDC1-CTF chimera enables Nanobody coating of a substantial proportion (6-20%, depending on the analytical approach) of CCM sEVs in the secretome in the absence of serum.

### Coating of sEV with anti-EGFR Nb via SDC1-CTF fusion improves sEV uptake by EGFR-expressing recipient cells

To be able to reliably quantify sEV uptake (binding and/or internalization), we aimed to load CCM sEVs with Nanoluciferase (NL): a small, highly sensitive and stable luciferase derived from deep-sea shrimp (Hall et al. 2012). We tested two options, loading with NL-syntenin or NL-CD63 fusion proteins. While both were efficiently loaded in sEVs, we observed cleavage products for NL-CD63 transfectants in the CCM sEVs (Fig. S6A). We therefore proceeded with NL-syntenin loaded CCM sEVs in further experiments. After optimizing NL substrate (membrane permeant Furimazine) concentration (Fig. S6B), we incubated Panc-1 cells for 4 h with different doses of CCM sEVs originating from NL-syntenin transfected HEK293 cells. We observed luminescence signals in recipient cells that increased proportionally to doses (doses of CCM sEVs were expressed as number of producing cells per single recipient cell at the time of the plating, p/r) before reaching a plateau (Fig. 5A). As this plateau was still in the linear range of the luminescence detection of the instrument, we could conclude that uptake was saturable at doses of ∼100 p/r. Encouraged by these initial results, we decided to extend the coating of our EVs with Nanobody targeting EGFR, a cancer promoting receptor expressed on Panc-1 cells (Fig. 5C, inset), we thus generated EGFR Nb SDC1-CTF clones (referred to as cEGFR) to start from homogeneous producing cells. We thus prepared and characterized by microscopy (Fig. 3A) and Western blot (Fig. 5B, Fig. S7), HEK293 clones secreting CCM sEVs coated with anti-EGFR Nb. We observed, according to super-resolution microscopy (Fig. S6C), that 19% of tetraspanin-positive cEGFR CCM sEVs were also Nb positive.

**Figure 5.**
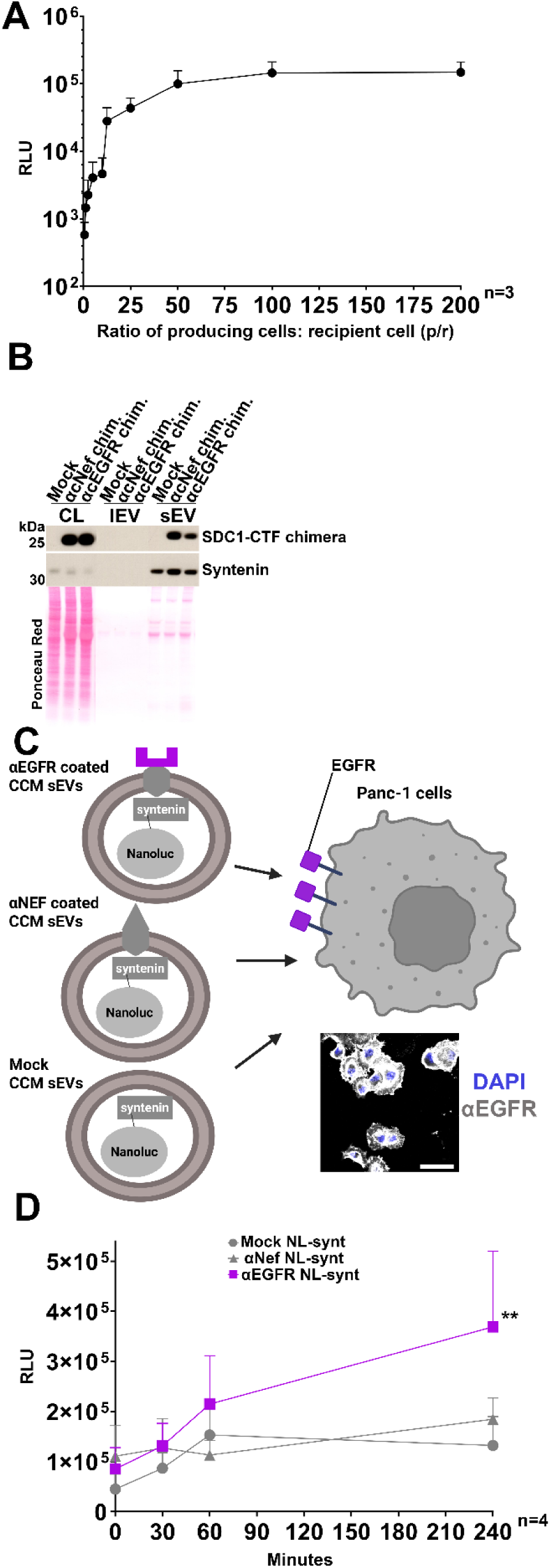
Coating of sEVs with anti-EGFR Nb improves uptake by EGFR-expressing cells. **(A)** Dose-response curve illustrating the uptake of non-coated CCM sEVs loaded with Nanoluciferase (NL)-syntenin. Panc-1 recipient cells were incubated for 4 h with increasing doses of CCM sEVs. (Y-axis) Luminescence signals (relative light units, RLU) were detected 5 min after washing Panc-1 cells, using the membrane permeant substrate Furimazine. (X-axis) Doses are expressed as the number of plated producing HEK293 cells per single plated Panc-1 recipient cell (p/r). Note the saturation of uptake from 100 p/r on. **(B)** Representative Western blot of the cell lysates (CL), large EV (lEV) and small EV (sEV) fractions (obtained by dUC) of cNef or cEGFR cells cultured in the absence of serum. Note that both chimeras are efficiently sorted into sEVs. Syntenin is used as sEV marker. Mock refers to HEK293 cells stably transfected with an empty vector. CL corresponds to 20.000 cells. EVs were collected from the conditioned media of 3.6 x 10^6^ cells. Ponceau red was used as loading and transfer control. All samples were loaded on the same membrane. For uncropped blot see Fig. S7. **(C)** Schematic representation of CCM sEVs coated with different Nb and loaded with NL-syntenin used for uptake experiments. Panc-1 cells were incubated with CCM sEVs under saturating doses (200:1 p/r). **Inset:** Representative confocal micrographs of Panc-1 cells illustrating their EGFR expression (grey). Nuclei are stained with DAPI (blue). Scale bar is 50 µm. **(D)** Time course of luminescence signals (y-axis, RLU) detected in Panc-1 cells following incubation with the different CCM sEVs as shown in **C**. Note that anti-EGFR CCM sEVs exhibit a significant increase in uptake compared to non-coated or anti-Nef CCM sEVs. Bars represent mean values + SEM. Statistical significance was determined by comparing the area under the curve using one-way ANOVA test with Tukey’s correction. Related to **Fig. S6 and S7**.

We investigated whether anti-EGFR sEV-coating improves uptake of NL-syntenin loaded sEVs compared to anti-Nef coated and control sEVs (Fig. 5C). We generated the respective NL-syntenin-loaded CCM sEVs (Fig. S6D) and confirmed that the total NL-syntenin signals were roughly equivalent for the different coatings (Fig. S6E). Then, we measured the uptake by Panc-1 cells over different periods of time at saturating dose of 200 p/r (Fig. 5D). Anti-EGFR coated sEVs were on average taken up three times better than non-coated or anti-Nef-coated sEVs after 240 min (Fig. 5D).

Taken together, these data indicate that NL-syntenin sEV uptake is saturable but can be increased in EGFR expressing cells when sEVs are coated with anti-EGFR Nb. Indirectly, these results also indicate that the present strategy and CCM sEVs preserve the activity of Nbs.

## Discussion

The field of EV therapeutics currently faces many challenges, including the design of reliable approaches improving EV targeting to diseased cells. In this study, we developed and validated a novel strategy to coat the surface of small extracellular vesicles (sEVs) using a chimeric protein composed of the cytosolic and transmembrane domain of the sEV marker SDC1 (Baietti et al. 2012; Castro-Cruz et al. 2023), fused to a juxta-membrane protease resistant peptide of the immune receptor CD4. We used different quantification approaches, including single EV fluorescence analysis and bioluminescence assays to show that this chimera can enable efficient sEV sorting of functional Nbs and improve syntenin-loaded sEV uptake.

Interestingly, our chimera appears as efficient as the previously described PTGFRN targeting strategy (Dooley et al. 2021). However, in our experiments, the PTGFRN pmNef produces high–molecular weight bands in Western blot of lysates and sEVs (Fig. 1C, 2A, S2, S3). We excluded this could be explained by aberrant processing that may occur in transient transfection experiments. The presence of such bands may correspond to SDS-insoluble aggregates and raises concerns in term of quality control. Aggregates are usually toxic (Bigi et al. 2023), which may raise safety issues. Although this would need to be experimentally addressed, another potential advantage of the SDC1-CTF strategy over the PTGFRN strategy is its small size (Fig. 1B), which may potentially improve membrane fusion and sEV internal cargo delivery.

Noteworthy, when comparing the secretomes of control cells with those of cNef, we observed no change in EV size or concentration using NTA or MRPS-approaches (Fig. 2B and 3C). One could argue that this may be related to the low percentage of chimera EVs (6-20%) and/or the suboptimal sensitivity of the two methods. Yet, in single EV analysis we also observed no evidence for shifts in concentration or size (Fig. 4C). This indicates that our strategy mainly comes with a change of sEV composition, with little if any impact on sEV size or concentration. Of note, the positivity 6-20% of the chimera EVs is comparable with other commonly used strategies for EV surface engineering such as for example PTGFRN or Lamp2b (Gupta et al. 2021). One aspect to consider for future studies is the knock-out of the endogenous SDC1, which may improve the sEV loading of chimera due to possible competition of the endogenous SDC1 with the chimera.

Different ‘single EV’ analytical approaches provided different results when quantifying the percentage of Nb-coated sEVs. The relatively low percentage observed in ARC^®^ and CytoFLEX Nano^®^ compared to ONI^®^ may be due to the fact that these approaches measure total particles that do not necessarily correspond to EVs. Another reason why the CytoFLEX Nano^®^ gave solely ∼6% may come from the need of a purification step to eliminate free labelled antibodies. This step of purification could be omitted with ARC giving ∼10% of Nb-coated EVs. The relatively high percentage observed in ONI microscopy (20%) might be the result of the double selection associated to the approach: (i) the capture was based on phosphatidylserine sEV composition and (ii) solely EVs that were positive for CD9, CD63 or CD81 (and at least 4 positive signals) were taken into consideration. Mock positivity observed in ONI likely reflects background binding and sensitivity limits of the technology rather than true nanobody decoration. Obviously, the physics behind the 3 instruments is very different which could equally impact the data and explain the differences. Errors in estimation are also expected to arise from differences in fluorophore performance. Alexa 647 was used for ONI^®^, while Alexa 488 was used for ARC^®^ and CytoFLEX Nano^®^. The comparison of these two fluorophores is not straightforward, as Alexa 488 is not optimal for ONI^®^ due to insufficient blinking, while Alexa 647 has solubility issues and is therefore not optimal for in solution approaches like Nano flow and MRPS (Linde et al. 2011; Dempsey et al. 2011). It would also be interesting to measure the amount of Nbs on the sEV surface by using fluorescent proteins fused to the SDC1-CTF chimera or by using non-antibody based approach, as antibodies have been recently suspected to underestimate the overall sEV display (Mitrut et al. 2024).

To quantitatively measure sEV uptake we opted for the Nanoluciferase (NL), as this luciferase is highly stable and sensitive (Hall et al. 2012). NL was loaded into sEVs by fusion to syntenin. As we previously established that syntenin directly interacts with SDC1-CTF (Baietti et al. 2012; Castro-Cruz et al. 2023), concurrent loading of both Nb and NL into the same sEV subpopulation is highly probable. Interestingly, we clearly observed a dose-dependent but saturable uptake of NL when incubating NL-syntenin loaded CCM sEVs with Panc-1 cells for 4 hours (Fig. 5A). It may be noted that saturation is not due to NL or sEV depletion, as the overall uptake remains low considering the total dose. Indeed, uptake never reaches 1% of the total administered NL-luminescent signal even after 4 hours (compare Fig. 5D and Fig. S6E). To our knowledge, only a few studies addressed in a quantitative manner the efficiency of EV uptake *in vitro* (Bonsergent et al. 2021; Hirose et al. 2022; Toribio et al. 2019), two of them reporting low percentages of EV uptake (Bonsergent et al. 2021; Hirose et al. 2022). Thus, limited uptake is certainly not restricted to NL-syntenin EVs, and clarifying whether such limited uptake is a general feature would require additional studies. Irrespective of these considerations, we could objectivate the benefit of Nb coating because our measurements indicated that anti-EGFR coated sEVs underwent three times more uptake than non-coated or anti-Nef-coated sEVs (Fig. 5D and S6D). Assuming the threefold increase in uptake pertains to the only 10-20 percent of NL-containing sEVs expressing Nb, anti-EGFR coating may thus increase sEV uptake by Panc-1 cells by nearly one order of magnitude. Likely, anti-EGFR supports an interesting receptor-mediated way of entry for sEVs.

In conclusion, the SDC1-CTF chimera is a useful genetic tool for engineering customized coated sEVs. It offers high sorting efficiency, presents no unexpected features and allows functional targeting. Compared to previously described strategies, including PTGFRN-based scaffolds and earlier syndecan-derived constructs, our approach provides a compact and protease-resistant design that ensures nanobody presentation. Pertinent perspectives include (i) the development of more elaborated technology that would allow easy purification of Nb-coated sEVs and (ii) testing the pharmacokinetics and pharmacodynamics of such vehicles *in vivo*. Also, it would be interesting to test the efficiency of our SDC1-CTF chimera approach to display poorly immunogenic peptides other than Nb (Rossotti et al. 2022) at the surface of sEVs.

## Data availability statement

The data generated in this study are available from the corresponding author upon request.

## Acknowledgements

The authors thank JB Vasquez Spectradyne^®^, C. Walentynowicz Analis^®^, P. Meuwissen Analis^®^, N. Nevo and the EV Curie Coretech for technical advice. This research project was funded by Kom op tegen Kanker (Stand up to Cancer), the Flemish cancer society (projectID: 12367) and the internal funds of the KU Leuven (C14/20/105). L.H. is a PhD Fellow of the Research Foundation – Flanders (FWO, 1SE2422N).

## Author contribution

P.Z., P.C., G.D. and L.H. conceived and designed the study. L.H., P.Z. and G.D. wrote the manuscript; L.H., S.M. and R.G performed experiments and analyzed the data. G.D., E.D.B., P.Z., and L.H. provided critical suggestions and revised the manuscript. All authors read and approved the final manuscript.

