## Supplementary information for "A syndecan-based genetic approach to coat the surface of small extracellular vesicles with Nanobodies"

### Supplementary figures

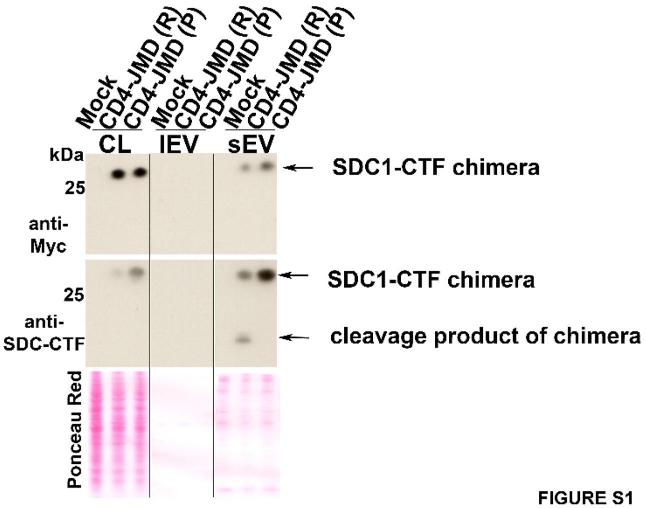


FIGURE S1

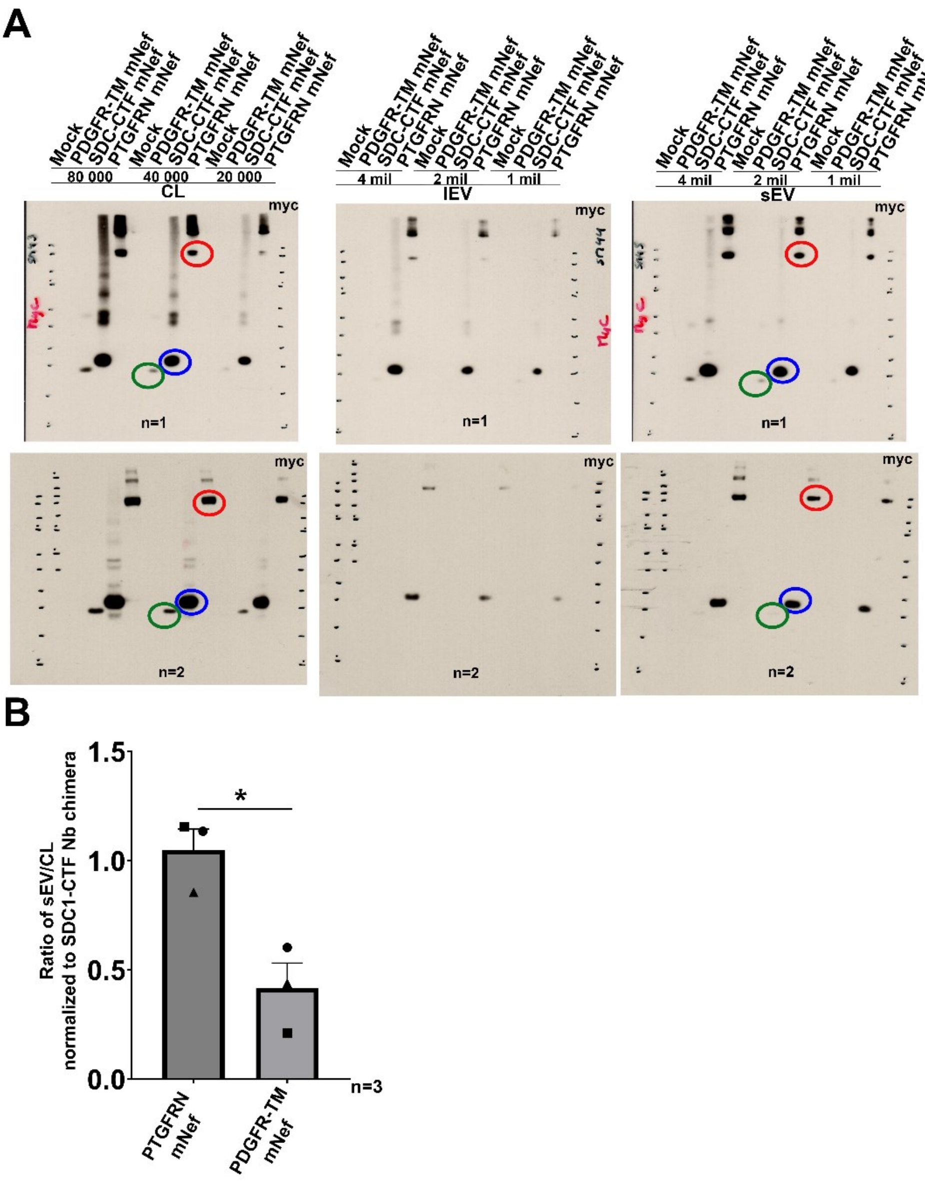


FIGURE S2

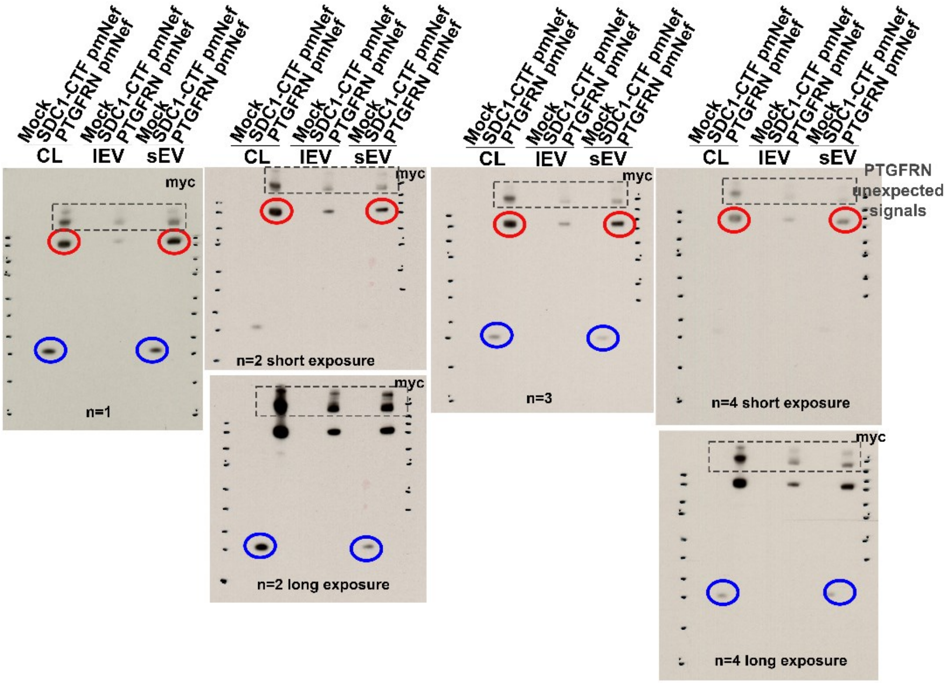


FIGURE S3

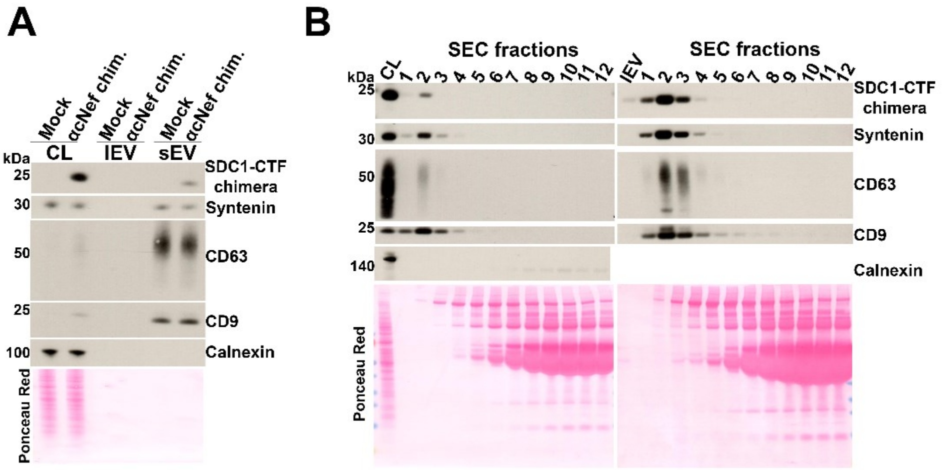


FIGURE S4

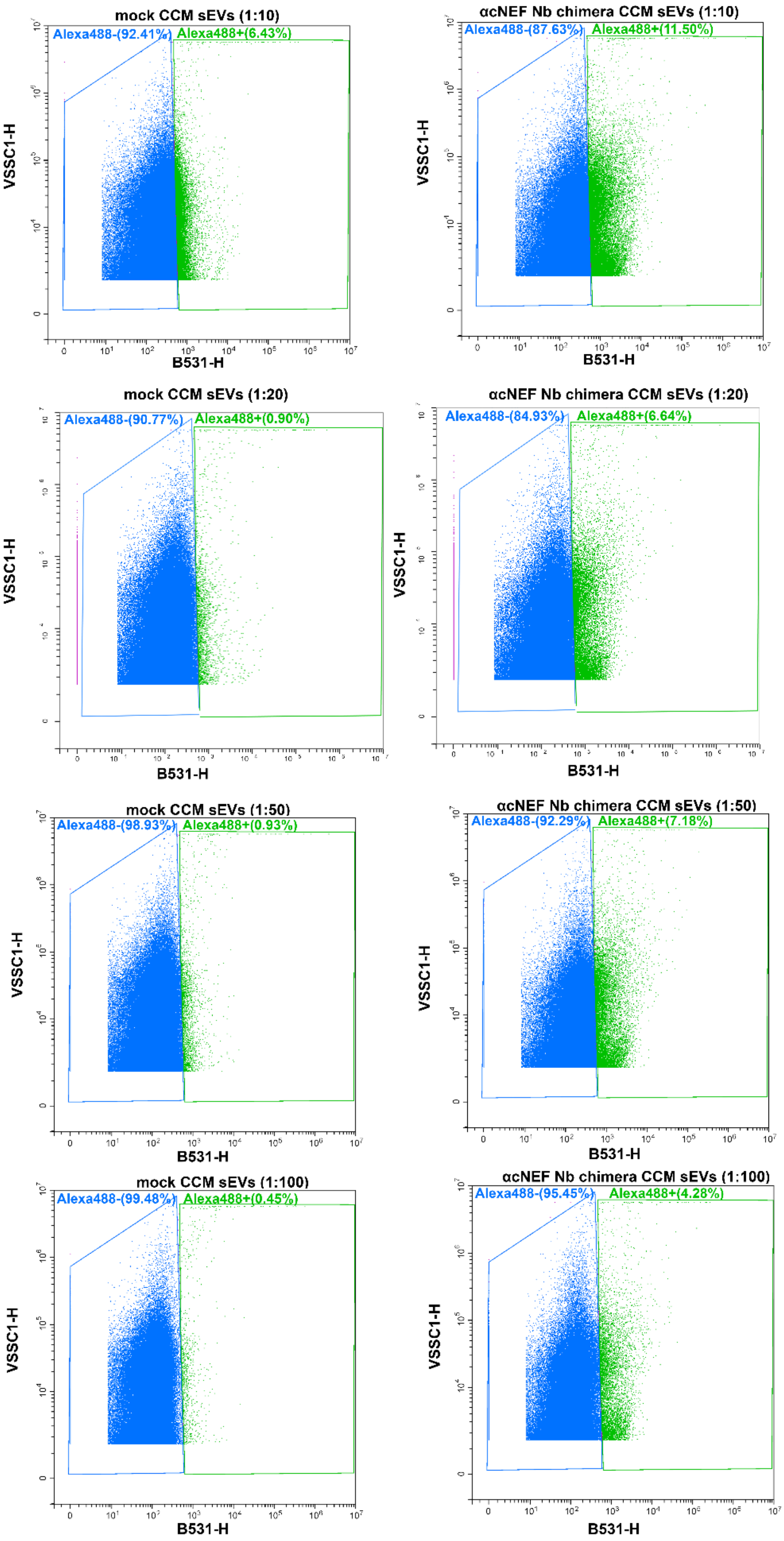
mock CCM sEVs (1:10)
Alexa488-(92.41%)
Alexa488+(6.43%)
VSSC1-H
B531-H
αcNEF Nb chimera CCM sEVs (1:10)
Alexa488-(87.63%)
Alexa488+(11.50%)
VSSC1-H
B531-H
mock CCM sEVs (1:20)
Alexa488-(90.77%)
Alexa488+(0.90%)
VSSC1-H
B531-H
αcNEF Nb chimera CCM sEVs (1:20)
Alexa488-(84.93%)
Alexa488+(6.64%)
VSSC1-H
B531-H
mock CCM sEVs (1:50)
Alexa488-(98.93%)
Alexa488+(0.93%)
VSSC1-H
B531-H
αcNEF Nb chimera CCM sEVs (1:50)
Alexa488-(92.29%)
Alexa488+(7.18%)
VSSC1-H
B531-H
mock CCM sEVs (1:100)
Alexa488-(99.48%)
Alexa488+(0.45%)
VSSC1-H
B531-H
αcNEF Nb chimera CCM sEVs (1:100)
Alexa488-(95.45%)
Alexa488+(4.28%)
VSSC1-H
B531-H

FIGURE S5

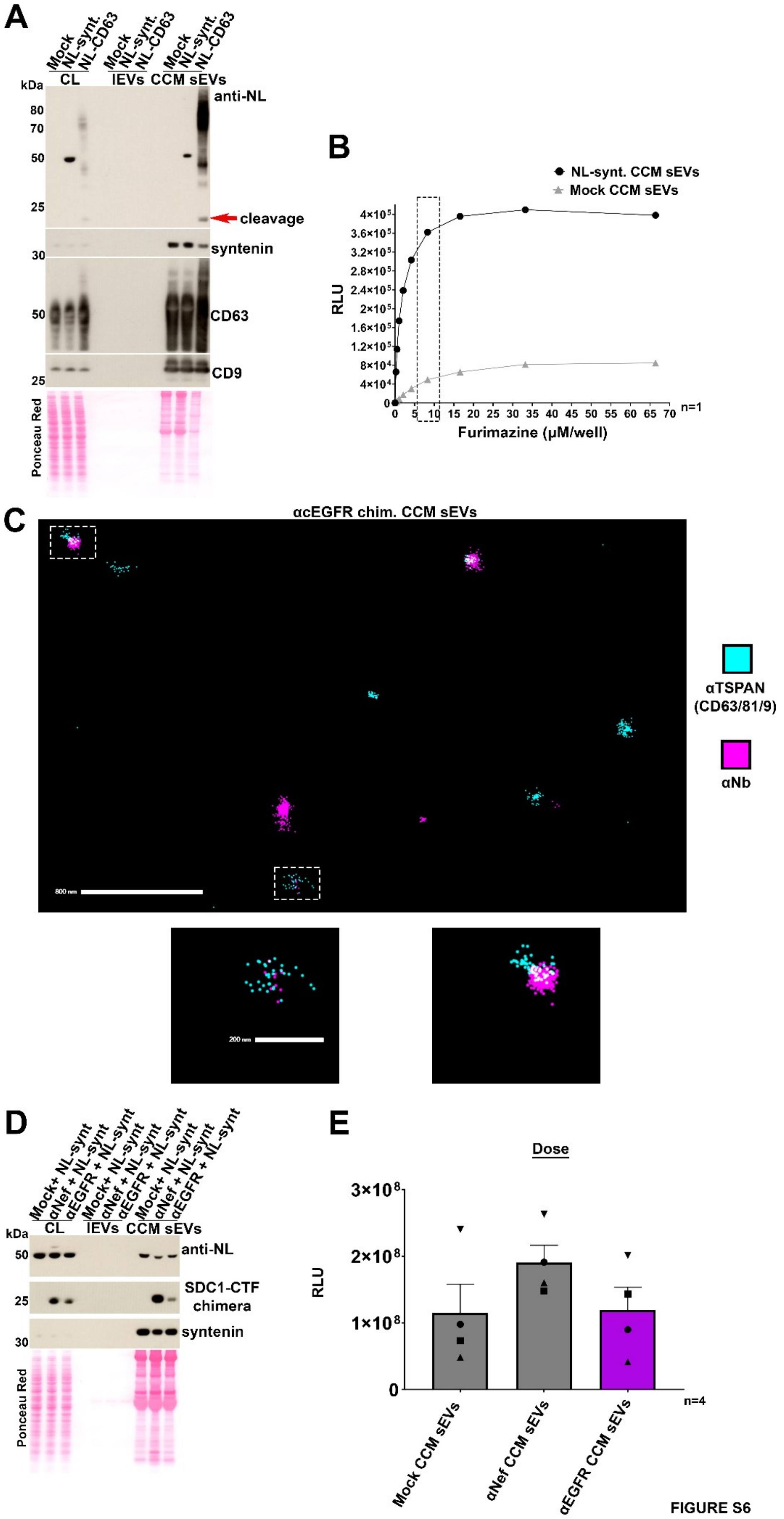
A
Mock
NL-synt.
NL-CD63
CL
lEVs
CCM sEVs
kDa
80
70
50
25
30
anti-NL
cleavage
syntenin
CD63
CD9
Ponceau Red
B
NL-synt. CCM sEVs
Mock CCM sEVs
RLU
Furimazine (µM/well)
n=1
C
αcEGFR chim. CCM sEVs
αTSPAN (CD63/81/9)
αNb
800 nm
200 nm
D
Mock+ NL-synt
αNef + NL-synt
αEGFR + NL-synt
SDC1-CTF chimera
E
Dose
3×10⁸
2×10⁸
1×10⁸
0
n=4
Mock CCM sEVs
αNef CCM sEVs
αEGFR CCM sEVs

FIGURE S6

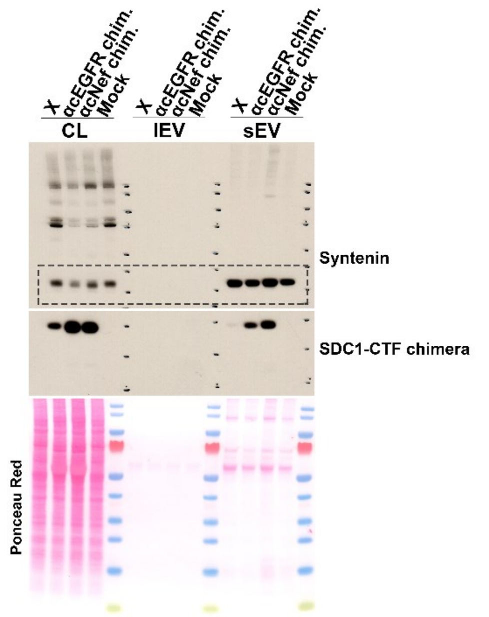


FIGURE S7

### Legends Supplementary figures

**Figure S1. Impact of the CD4 juxta-membrane sequence on the cleavage of SDC1-CTF chimera.** Illustrative Western blot of cell lysates (CL) and EVs (fractionated by dUC) from HEK293 cells transiently expressing myc-tagged Nef Nb chimera containing a CD4 juxta-membrane domain with an arginine (CD4-JMD (R)) or a proline (CD4-JMD (P)). Large EV (lEV) and small EV (sEV) fractions correspond to the 10K and 100K pellets, respectively. Blots were probed with anti-myc antibodies (upper panel), an epitope present at the N-terminus of the chimera, or anti-SDC1 antibodies (middle panel) recognizing the C-terminus of the chimera. CL correspond to 20.000 cells. Secretomes were collected from the conditioned media of 3.6 x $10^6$ cells. Red Ponceau (lower panel) was used as loading control. Images are from the same blot. Note the presence of a cleavage product in the sEV fraction of CD4-JMD (R) transfectants. Related to **Fig. 1A.**

**Figure S2: PDGFR-TM chimera is not efficient at sorting Nanobodies to extracellular vesicles. (A)** Uncropped Western blots of HEK293 cell lysates (CL), and large and small EV (lEV, sEV) from cells transiently transfected with various constructs as indicated. Mock refers to cells transfected with an empty vector. For CL, numbers refer to cell amounts. For lEV and sEV fractions, numbers correspond to the amount of producing cells. Anti-myc antibodies were used to compare the distribution of SDC1-CTF, PTGFRN, and PDGFR-TM myc-tagged Nef Nbs, as indicated. Note the unexpected signals detected in

SDC1-CTF and PTGFRN transfectants. Circles indicate areas used for quantification of sorting efficiencies. Blue for SDC1-CTF, red for PTGFRN and green for PDGFR-TM chimeras. **(B)** Histogram illustrating that PDGFR-TM chimera is less efficient at sorting Nbs to sEVs than SDC1-CTF chimera. Anti-myc signals obtained in Western blot for sEV fractions and CL, were normalized to SDC1-CTF chimera signals in their respective fractions and expressed as a mean ratio sEV/CL from 3 independent experiments. Bars represent mean values + SEM. Student's t-test was applied to assess statistical significance. Related to **Fig. 1B-C.**

**Figure S3. Unexpected signals remain present in cells stably overexpressing the PTGFRN chimera.** Uncropped Western blots of HEK293 cell lysates (CL), and large and small EVs (lEV, sEV), from cells stably overexpressing SDC1-CTF and PTGFRN chimeras. Mock refers to cells transfected with an empty vector. CL correspond to 20.000 cells. EVs were collected from the conditioned media of 3.6 x $10^6$ cells. Circles indicate areas used for the quantification of sorting efficiencies of SDC1-CTF (blue) and PTGFRN (red) chimeras. Related to **Fig. 2A.**

**Figure S4. dUC characterization of cNef and additional SEC blots. (A)** Illustrative Western blot of EVs fractionated by dUC (lEVs and sEVs pellets, as indicated), along with the corresponding cell lysates (CL) from the HEK293 clone stably overexpressing the SDC1-CTF Nef Nb chimera. Mock refers to HEK293 cells stably transfected with an empty vector. Various EV marker proteins used as positive controls and calnexin used as negative control, as indicated on the right. Ponceau red was used as loading and transfer control. Note that the chimera is detected in CL and sEVs but is absent from lEVs. **(B)** Western blots showing two extra biological repeats of sEV fractionation by SEC experiments as in **Fig. 3D**. Related to **Fig. 3** and **Fig. 5.**

**Figure S5. Nano flow analysis data.** Dot plots of CCM sEVs from mock-transfected (left) or anti-Nef CCM sEVs (right). CCM sEVs were stained with Alexa488-conjugated anti-Nb antibody, using different dilutions. The gating strategy was used to separate the bulk of dense signal (blue) of the Mock CCM sEVs treated with 1:20 antibody dilution (concentration used throughout this study for single-EV detection) and transposed to all the other samples. Percentages of Nb negative (blue) and positive (green) particles are indicated. Related to **Fig. 3.**

**Figure S6. Characterization of Nanoluciferase-syntenin loaded CCM sEVs by Western blot and luminescence. Single-vesicle characterization of anti-EGFR CCM sEVs. (A)** Representative Western blots of the cell lysates (CL) and secretomes (lEV and CCM sEVs) of HEK293 cells transiently overexpressing Nanoluciferase-syntenin (NL-synt) or Nanoluciferase-CD63 (NL-CD63), as indicated. Mock refers to HEK293 cells transiently transfected with an empty vector. Blots were probed with anti-NL (upper panel) or sEV marker antibodies, as indicated on the right. Ponceau red was used as loading and transfer control. Note that both NL-synt and NL-CD63 are efficiently sorted to CCM sEVs. Also note the low-molecular-weight signal (red arrow) in the NL-CD63 CCM sEVs suggesting cleavage of the NL part from the NL-CD63. CL correspond to 20.000 cells. Secretomes were collected from the conditioned media of 4.8 x $10^6$ cells. **(B)** Dose-response curve illustrating the luminescence signals (relative light units, RLU, Y-axis) obtained with NL-synt loaded (black) or mock CCM sEVs (grey) at increasing concentrations (µM/well, X-axis) of the Nanoluciferase substrate, Furimazine. From there, we decided to set the Furimazine concentration at 8.3 µM (indicated in dashed frame). **(C)** Representative micrographs of anti-EGFR CCM sEVs characterized by ONI® microscopy. Anti-tetraspanin (CD9, CD63, CD81) signals are in cyan while anti-Nb signals are in magenta. Scale bars correspond to 800 nm for field view and 200 nm for inserts. **(D)** Illustrative Western blot of CCM sEVs prepared from the different SDC1-CTF chimera clones also overexpressing NL-syntenin, as indicated on the top. Blots were probed with anti-NL antibodies, anti-SDC1 antibodies recognizing the intracellular domain of SDC1, and anti-syntenin antibodies recognizing the endogenous syntenin. Ponceau red was used as loading and transfer control. CL corresponds to 20.000 cells. The secretome was prepared from conditioned media of 2.4 x $10^6$ cells. **(E)** Bar graph showing the luminescence signals (RLU, Y-axis) from NL-syntenin loaded CCM sEVs obtained from cells stably expressing the different SDC1-CTF chimeras (as indicated on the X-axis). Individual points represent individual doses administered to Panc-1 cells in 4 independent biological repeats. Bars show mean values + SEM. Related to **Fig. 5.**

**Figure S7. Uncropped blot of cNef and cEGFR cells.** Uncropped western blot of the cell lysates (CL), large EV (lEV) and small EV (sEV) fractions (obtained by dUC) of cNef and cEGFR HEK293 cells. Dashed area shows cropped part of the anti-syntenin blot used in **Fig. 5.**
